# The One Click Wonder: a retrained automated segmentation pipeline that enables quantitative and modular analysis of *C. elegans* embryos

**DOI:** 10.64898/2026.01.21.700865

**Authors:** Palmer Carlyn Bassett, Tobias Evan Verheijen, Angelo Luiz Angonezi, Aude Andriollo, Sebastien Herbert, Gregory Roth, Jeffrey A. Chao, Susan E. Mango

**Author notes:** Equal contribution (Palmer Carlyn Bassett, Tobias Evan Verheijen).

## Abstract

High-throughput approaches have transformed the study of gene regulation by enabling quantitative, genome-scale analyses in both genomics and imaging. However, applying these methods to intact organisms remains challenging, particularly for high-throughput, 3D imaging. In Caenorhabditis *elegans*, generalist segmentation models often perform poorly due to rapid changes in nuclear size, shape, and density. To overcome this obstacle, we developed **One Click Wonder (OCW)**, an automated pipeline that pairs a retrained Cellpose model with stage-specific parameter selection to deliver accurate, high-throughput segmentation of embryos. We further introduce the **Biological Annotation and Association Mapper (BAAM)**, which integrates segmentation with spot detection, to enable single-cell quantitation. Applied to the pioneer factor *pha-4/FoxA*, this pipeline revealed distinct cell populations with an eight-fold range in transcriptional burst frequency. These findings demonstrate that OCW and BAAM provide a modular, scalable pipeline for quantitative, single-cell analysis of gene expression in *C. elegans* embryos.

## Background

High-throughput analysis of biological traits has revolutionized the scale and depth of our understanding of biology. Genomics methods such as RNAseq, ChIP and Hi-C have captured genome-wide information for transcription, regulation and 3D chromosome organization. More recently, advances in high-throughput image analysis are transforming our ability to quantify complex cellular dynamics for the spatial and temporal distributions of proteins and nucleic acids. Nevertheless, quantitative analysis at single-cell resolution in multicellular organisms continue to present a significant computational bottleneck. Specifically, segmentation of three-dimensional microscopy images remains challenging for small, dynamic systems like *Caenorhabditis elegans* embryos. Traditional segmentation models and deep-learning based tools often underperform on images from biological systems such as *C. elegans*, which exhibit rapidly changing cellular architecture, tight cell packing, and variable imaging conditions^1,2^. Compounding this problem, experimental pipelines often produce heterogeneous image formats and resolutions, which demand computational solutions that are not only robust, but also modular and adaptable for diverse analysis workflows.

Classical image segmentation approaches, such as watershed-based methods have long served as a foundation for quantifying cellular features in microscopy data^3–5^. However, these classical methods often demonstrate poor segmentation quality, particularly in systems like developing embryos where cell size, shape and spacing vary over time. These methods require extensive manual parameter and threshold tuning, which are sensitive to noise and signal variability, making reproducibility across experiments, datasets and researchers difficult. Recent advances in deep learning-based segmentation, with tools such as Cellpose, StarDist, SAM and Omnipose have markedly improved performance on many benchmark datasets^1,6–8^. However, these models underperform when applied to biological systems outside their training domain, particularly with noisy, blurred or under-sampled 3D images^1^. Domain-specific retraining can significantly improve performance, as demonstrated in *C. elegans* germline nuclei segmentation done by Padovani et al., 2022^9^, but such improvements do not generalize to embryonic datasets, which undergo rapid morphological changes during development. As a result, even state-of-the-art tools require substantial customization to reliably perform in dynamic, non-canonical imaging contexts, underscoring the need for robust modular pipelines.

To address these downfalls, we developed a machine learning-based image processing pipeline, designed to enable fast, reproducible, and accurate three-dimensional segmentation of *C. elegans* embryos. Affectionately named the *One Click Wonder* (OCW), the pipeline incorporates a retrained Cellpose model, supports multiple image formats, and enables robust segmentation of embryonic nuclei. Crucially, it accounts for the dynamic morphological changes that occur during embryogenesis through a stage-specific neural network parameter selector, which applies optimized segmentation parameters tailored to each developmental stage.

To demonstrate its utility, we applied the One Click Wonder to a biological case study: transcriptional bursting during *C. elegans* embryogenesis. Transcriptional bursting is a stochastic process in which genes switch between active and inactive transcriptional states, episodically producing RNA^10^. Bursting influences cell-to-cell variability in gene expression and plays a key role in development. We focused on *pha-4*/FoxA, a pioneer transcription factor essential for the pharyngeal organogenesis in *C. elegans*^11–15^. Using RNA smFISH and established dot-detection methods^16^, we demonstrate the modular capabilities of the OCW pipeline to quantify nascent *pha-4* transcripts and applied a two-state model to infer bursting kinetics.

This study establishes the One Click Wonder as a robust and adaptable segmentation tool for developmental imaging and demonstrates its capacity to support detailed, quantitative analyses of transcription at single-cell resolution. Our pipeline, in combination with dot-detection and modeling methods, revealed two transcriptionally distinct cell populations in developing embryo, in which *pha-4* exhibits differential bursting frequences. We note that the OCW is not restricted to RNA analysis, but can be applied for other analyses such as protein stains or DNA FISH.

## Results

We developed the One Click Wonder (OCW), an automated approach for segmenting nuclei and embryos from individual images or image sequences. This method integrates image preprocessing, machine-learning-based segmentation, and image classification into a single workflow optimized for *C. elegans* imaging. We then highlight how this method can be used in combination with other computational outputs such as TrackMate and give a specific example of how this method can be used to reveal differential transcriptional dynamics at a single-cell level.

We created and optimized an automated pipeline for single-cell analysis of *C. elegans* embryos. The performance of the Cellpose *cyto2* model was evaluated out of the box to determine its suitability for 3D segmentation of *C. elegans* embryo samples, but we found the pretrained *cyto2* model struggled to segment nuclei accurately. In early-stage embryos (0-40 CS), the model frequently over-segmented nuclei, where a single nucleus was apparent in the raw image, Cellpose generated multiple, smaller, and incomplete segmentation labels (Fig. 1A). *cyto2* also frequently under-segmented nuclei, particularly apparent in mid-stage embryos (41-100 CS), by merging adjacent nuclei or missing some nuclei altogether (Fig. 1B). Nuclei were often incompletely labeled by Cellpose *cyto2*, failing to identify nuclei that were packed closely (Fig. 1C). This is most evident in late-stage embryos (150+ CS) where cells are tightly packed within the embryo. These segmentation errors across developmental stages indicated that retraining the *cyto2* model was necessary to achieve reliable and suitable nuclear segmentation for *C. elegans* embryos.

**Figure 1:**
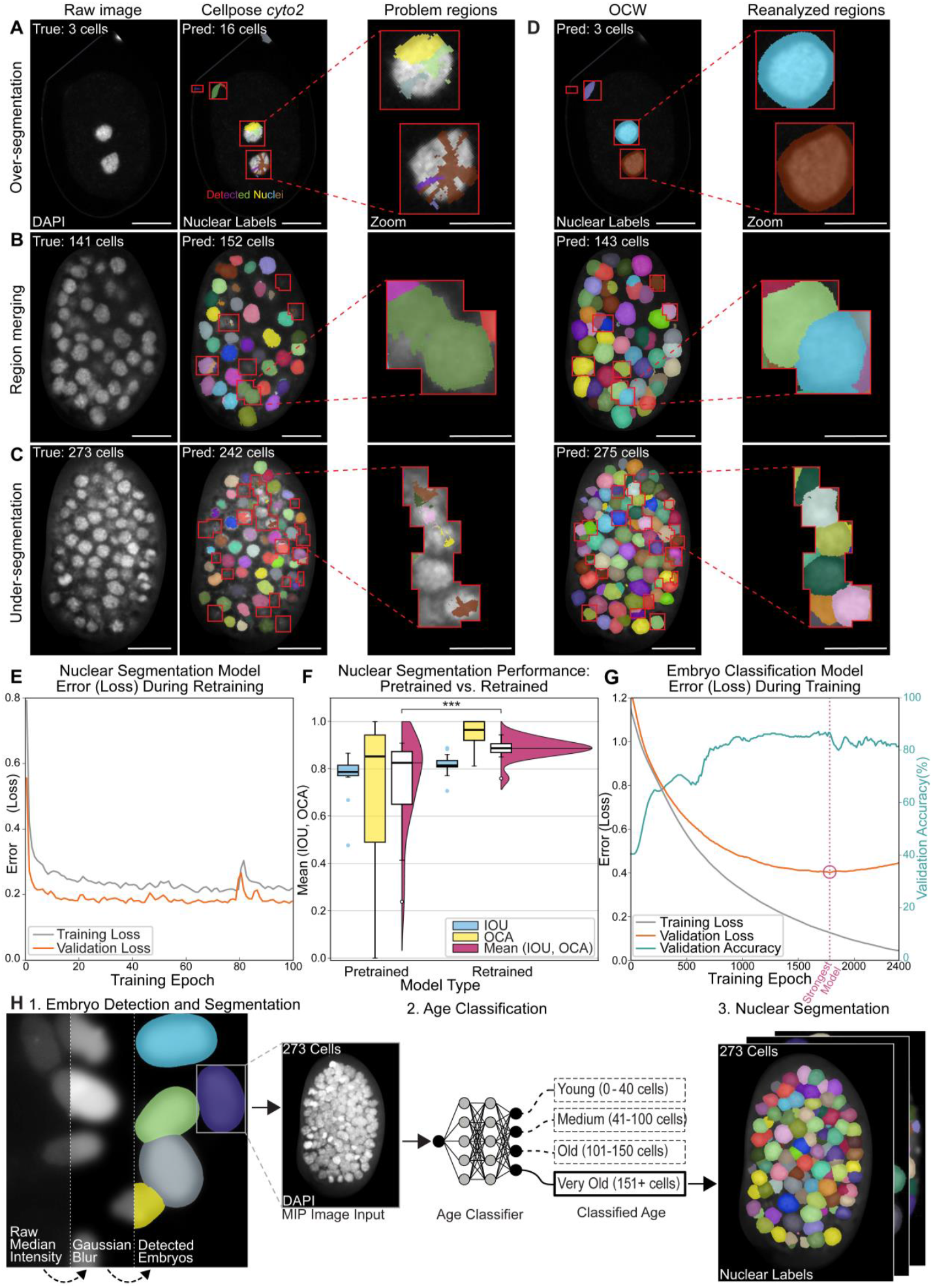
The One Click Wonder provides a clean, accurate and efficient method to segment nuclei and embryos from a single or sequence of image(s). A, B, & C: Segmentation problems with base Cellpose *cyto2* model. Representative images of DAPI staining of embryos (raw image) from three age ranges (A, early: B, mid-stage, C, late-stage), followed by the detected object labels output from the base Cellpose *cyto2* model with the best set of parameters. True ages are displayed in the top left corner of the images in the first column of raw images. The number of labels detected is displayed in the top left corner of the images in the second column, Cellpose *cyto2*. Red boxes demonstrate lackluster segmentation results with under-segmented, over-segmented or merged labels. A zoom of each problem region is provided. Scale bar 10 mm for DAPI and Detected Nuclei and 5mm for Zoom. D: OCW segmentation performance. The left panels display detected nuclear labels from retrained Cellpose *cyto2* model from the embryos shown in panel A. Red boxes correspond to the problem regions for the base *cyto2* model. The number of labels detected is given in the top left corner of the first column. Right panels show enlarged regions that were problematic in Cellpose *cyto2*, but corrected in OCW. E: Nuclear segmentation model error (loss) during retraining of the Cellpose *cyto2* model. Error (loss) is on the y-axis and training epochs is on the x-axis. Training loss is shown in grey while validation loss is shown in orange. F: Cellpose model quality demonstrated through the intersection over union (IOU) and Object Count Accuracy (OCA) metrics. The magenta regions with white boxes represent the combined IOU and OCA scores, computed by taking the mean of the two statistics. A paired t-test was performed with a ‘less’ alternative hypothesis on the combined scores which yields significance with a *p*-value less than 0.005, indicated with three asterisks. The blue and yellow box and whisker plots show the influence of retraining on the IOU and OCA scores. IOU and OCA scores (0-1) are on the y-axis and model type is on the x-axis. G: Embryo classification model error (loss) during training and validation accuracy for the training of our bespoke embryo age classifier. The training is run past the point of overfitting to find the lowest validation error (loss), this point is indicated by the “Strongest Model” dotted magenta line and circle. Error (loss) is on the left y-axis, validation accuracy is on the right y-axis, and training epochs are on the x-axis. Training loss is shown in grey, validation loss in orange and validation accuracy is displayed in teal. H: Schematic overview of OCW workflow. (1) A median projection of the raw image of a uniform background intensity channel is taken, and Gaussian blurring and re-contrasting is applied. Embryos are detected in 2D using Cellpose *cyto2*. (2) Age classification is performed by taking a DAPI maximum intensity projection and passing it to an age classifier. The age class is given as input to (3). (3) The retrained Cellpose *cyto2* model is run in 3D on embryos (embryo label broadcasted through full z-stack) using the parameters defined by the classified age group.

Following the recommendations outlined by Pachitatiu & Stringer^33^, we retrained the model on a curated dataset (see Cellpose Supplementary). After retraining, segmentation performance improved substantially, enabling accurate nuclear detection across all developmental stages, from embryos with 2 to 558 cells. The issues of over-segmentation, under-segmentation, and region merging observed in the base *cyto2* model were effectively resolved (Fig 1D). To assess model performance during training, we monitored the training and validation losses (Fig 1E). The progressive decline in loss values reflects convergence of the model parameters toward improved segmentation performance. To compare the base and retrained models quantitatively, we calculated intersection over union (IOU) and object count accuracy (OCA) metrics. The retrained model exhibits a higher mean IOU and OCA, along with a considerable reduction in variability (Fig. 1F). This result indicates improved spatial accuracy of the segmentation labels and more accurate object counts relative to the ground truth.

To optimize segmentation performance and speed up the analysis pipeline, we performed a grid search to identify the optimal parameter sets for distinct developmental stages. Embryos were categorized into four age groups: Young (0–40 cells), Medium (41–100 cells), Old (101–150 cells), and Very Old (150+ cells). To avoid manual parameter selection, we developed an embryo age classification model to automatically determine the age group of each embryo and thus assign the respective set of parameters to be used. Bounding boxes from the segmented embryos were used to extract maximum intensity projections of the DAPI channel, which were taken as input to a ResNet-like convolutional neural network (He et al., 2016)^34^. During training of the embryo age classification model, the model loss initially decreased on both the training and validation sets. However, as training progressed, overfitting was observed: while the training loss continued to decrease, the validation loss began to rise. The model corresponding to the lowest validation loss was selected as the final classifier (Fig. 1G).

These components were integrated into an automated pipeline that consists of three primary stages (Fig. 1H): (1) embryo detection and segmentation, (2) age classification, and (3) nuclear segmentation. In the first stage, preprocessing is applied to a diffuse background channel by computing a median projection, followed by Gaussian blurring and contrast normalization. The processed image is then passed to Cellpose for embryo segmentation. In the second stage, age classification is performed on the segmented embryos. In the third stage, the appropriate retrained Cellpose parameters—determined by the predicted age group—are applied to conduct full 3D nuclear segmentation. We have renamed this the *One Click Wonder* (OCW) pipeline for its ease and speed of use.

The OCW pipeline substantially reduces processing time compared to the previous manual pipeline. Previous segmentation pipelines required up to one week of hands-on analysis per dataset and the time-intensive and tedious setting of parameters. The OCW completes the entire segmentation process in under 30 minutes when executed in a parallel analysis format. When relying on GPU resources but not executed in a parallel analysis format, the processing time is still significantly reduced to about 2-3 hours (dataset of about 20 images).

To extend the utility of OCW and enable downstream spatial quantification, we developed BAAM (Biological Annotation and Association Mapper), a modular set of pipelines that integrates OCW outputs with the positional data from dot-detection tools (e.g., TrackMate)^16^, by mapping detected puncta onto their corresponding embryonic or nuclear regions (Fig. 2A). We first used BAAM to quantify nuclear-localized intronic RNA smFISH signal in developing embryos. Combining OCW-segmented nuclei with detected intronic dots enabled accurate assignment of transcription sites at single-cell resolution (Fig. 2B). In addition to counts, BAAM also extracts signal intensities (Fig. 2C), which allows us to quantify additional transcriptional qualities.

**Figure 2:**
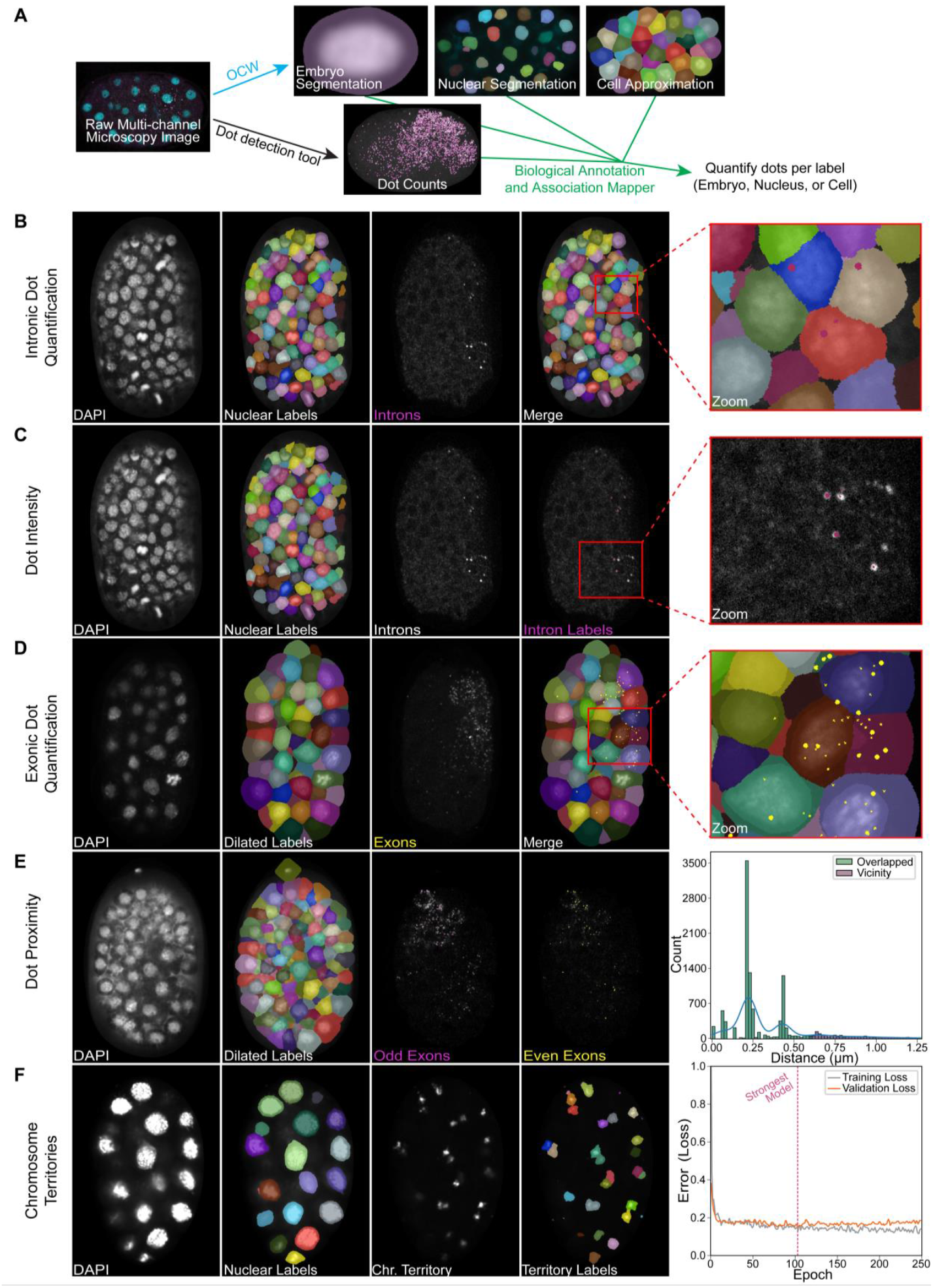
BAAM for nuclear, cytoplasmic and spatial RNA analyses. A: Schematic overview of the Biological Annotation and Association Mapper (BAAM). The general image processing workflow starts with raw multi-channel microscopy images. Top row: OCW is run which produces embryo, nuclear and cell approximation label images. Bottom row: Dot detection tools are run separately, producing dot-detection label images. These outputs are combined by the BAAM pipeline to quantify dots per label. B-F: OCW outputs used in combination with dot detection algorithms for smFISH dot detection (B-E) and chromosome territory segmentation methods (F). The first column shows the raw DAPI image, the second column shows the OCW output label image, the third column shows the externally-input dot detection result at a single z-slice or chromosome territory label image, and the fourth column shows the merged image (respective of row) and a zoomed image. For dot proximity analysis (E) a histogram of the distance in μm between exonic dots is shown as a representative example. Overlapped (<0.6 μm) and dots in the “Vicinity” (>0.6 μm) are shown as an example. For chromosome tracing, the chromosome segmentation model error (loss) during retraining of an additional Cellpose *cyto2* model is shown (F). Error (loss) is on the y-axis, epoch is on the x-axis, and training loss is shown in grey while validation loss is shown in orange.

For cytoplasmic exonic signal, we applied a difference-of-Gaussians detector alongside the OCW. Nuclear masks were dilated to approximate cell volumes, and BAAM mapped exonic positional data to these cell-proxy labels to estimate mature mRNA levels per cell (Fig. 2D).

We also designed BAAM to support spatial colocalization analysis across channels. As a test-case, we quantified distances between odd and even exon signals of a developmental gene, demonstrating rapid, automated assessment of signal colocalization within a single nucleus (Fig. 2E). BAAM for colocalization begins with the raw image, runs OCW, and has an integrated difference-of-Gaussians dot detection analysis.

Finally, we extended the OCW framework to segment chromosome territories. While not an explicit part of BAAM, we thought it important to include as it is a post-OCW processing modal. Developmental-stage specific parameters, inferred from nuclear counts, guided segmentation at different stages, enabling accurate territory identification across embryonic ages (Fig. 2F).

To test the utility of the OCW and BAAM, we applied it to a biological case study: *pha-4* gene expression during *C. elegans* embryogenesis. The gene *pha-4* is essential for pharyngeal organogenesis in *C. elegans*^35^. This case study serves two purposes: first, it demonstrates the utility of the OCW pipeline in combination with customized computational tools to quantitate nascent and mature RNA; and second, to characterize the transcriptional behavior of *pha-4*, a key developmental regulator. As a starting point, we confirmed that *pha-4* exhibits stochastic transcriptional behavior. RNA smFISH using intron- and exon-targeting probes (Fig. 3A) revealed cells with high mRNA abundance but no intronic signal, which is consistent with burst-like transcriptional dynamics.

**Figure 3.**
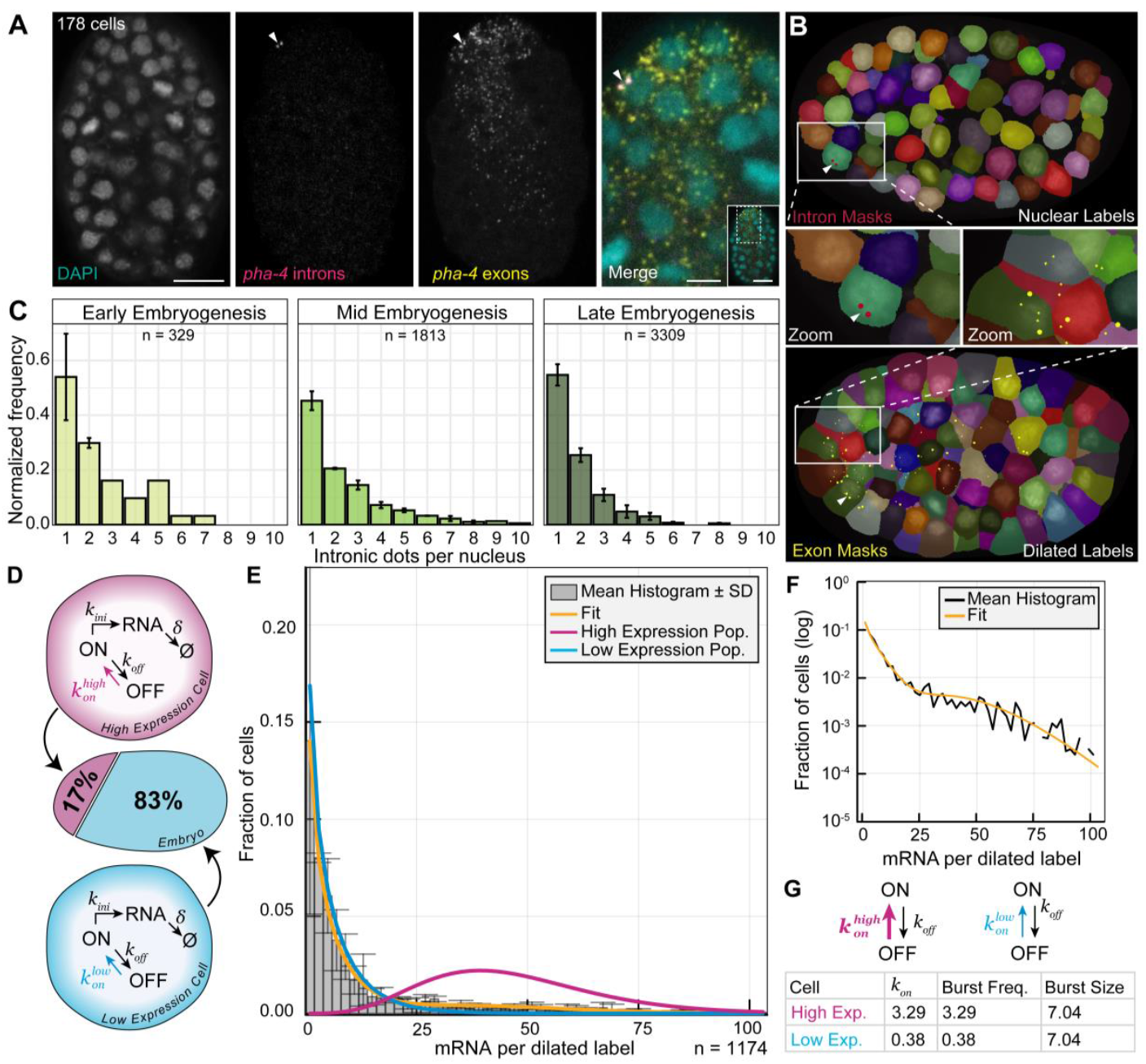
OCW with BAAM and mathematical modeling reveal two transcriptionally distinct *pha-4* populations in the embryo, with one exhibiting a burst frequency eight times higher than the other. A: Representative images of a 150-200 cell embryo with DAPI, *pha-4* RNA smFISH with probes targeting the introns, *pha-4* RNA smFISH with probes targeting the exons, and the zoomed merge image taken from the anterior portion of the embryo. Arrow heads point to transcription sites revealed by *pha-*4 intron probes. Scale bar 10 μm in DAPI and 5 µm in the Merge zoom. B: OCW and BAAM reveal the cell-specific location of RNA dots. Nuclear labels combined with TrackMate-detected intronic dots (top) and nuclear labels dilated by 40 pixels as a proxy for cell volume with TrackMate-detected exonic dots (bottom). C: Intronic dot counts per nuclear label for early (0-28 cells), mid (28-150 cells) and late (150-550) embryogenesis embryos. *N* counts represent the number of nuclei in each age-range. Frequencies are normalized. D: Schematic of the underlying assumptions for the two-population, two-state model. The bursting dynamics of each cell is described by a two-state model in which the promoter stochastically switches with rates *k*_*on*_ and *k*_*off*_ between an ON and an OFF state in which transcription can initiate with rate *k*_*init*_. RNAs are degraded at rate *δ*. The proportion of “high-*pha-4*-expression” cells is fixed to 17% of the embryo, as a 150-200 cell embryo has approximately 25 pharyngeal and intestinal precursor cells (Sulston et al., 1983)^35^. The “low-*pha-4*-expression” population is composed of the remainder of the cells (83% of the embryo). E: The best fit of the two-population, two-state model. Graph showing the distribution of mRNA per dilated nuclear label, with the x-axis representing the number of mRNA per dilated nuclear label and the y-axis indicating the fraction of cells. The observed mRNA counts are shown as a histogram. The yellow line shows the best-fit of the model. The blue and the pink lines show the model prediction of the mRNA distributions of the “low-*pha-4*-expression” and “high-*pha-4*-expression” populations, respectively. F: The two-population, two-state overall model fit on log-transformed fractions of cells to show relative fit in cells at the tail of the distribution. G: The two-population, two-state model reveals an eight-fold difference in burst frequency. Table depicts the *k*_*on*_ values, burst frequency, and burst size outputs of the model. The schematic is a visual representation of the *k*_*on*_ values in the table, with pink and blue representing the “high-*pha-4*-expression” and “low-*pha-4*-expression” cell types respectively.

To analyze *pha-4* transcription across embryogenesis, we required quantitative estimates of nascent RNA and mRNA per cell. Using the OCW, we performed nuclear segmentation and expanded the nuclear labels by 40 pixels to approximate cell boundaries (Fig. 3B). We ran TrackMate for dot-detection and used BAAM for assignment of transcription dots to their corresponding segmented nuclei and embryos. These segmentation-based assignments enabled an approximate per-cell quantification of nascent RNA and mRNA counts.

To investigate transcription on an age-specific level, we binned embryos by nuclear count into three groups: early-stage (0-28 cells), mid-stage (29-150 cells), and late-stage (150-500 cells). During the early stage, the embryo undergoes a few divisions and the initial pharyngeal precursor cells are specified. Mid-stage embryos undergo gastrulation and germ layer formation, while late-stage embryos undergo tissue differentiation and morphogenesis^36^. We found that early-stage embryos displayed the greatest variability in intronic dot number per nucleus, suggesting intron-containing RNA may accumulate in early-stage embryos more than late-stage embryos (Fig. 3C). Mid- and late-stage embryos generally showed 1-2 transcription dots per nucleus, which is the expected number of transcription dots for an actively transcribing diploid cell.

For transcriptional bursting analysis, we used the mRNA counts per cell derived from OCW and the exonic dot detection (Fig. 3B). Our analysis focused on embryos at the 150-200 CS, during which PHA-4 protein is thought to be actively produced^35^. At this stage, *pha-4* is expressed in two biologically distinct cell populations (Fig. 4D): a “high-expression” population comprising ∼25 pharyngeal and intestinal precursor cells (approximately 17% of the embryo), and a “low-expression” population representing the remaining 83% of cells.

We modeled each population independently using a standard two-state model of transcriptional bursting^37^, in which the promoter switches stochastically between an OFF state and an ON state, with rates *kon* and *koff* respectively. Transcription initiates from the ON state at rate *k*_*init*_ and mRNAs degrade at rate *δ* . The high- and low-expression populations were modeled with distinct parameter sets 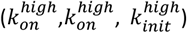 and 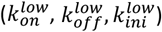.

We then asked whether the observed distribution of mRNA counts in 150–200 cell embryos could be explained under the hypothesis that the two populations differ in only one of these rates. Notably, the best fit was achieved when the *on* rate was the only differing parameter. This best fit model predicted an eight-fold higher *k*_*on*_ in the high-expression population compared to the low-expression one, corresponding to an eight-fold difference in burst frequency (Fig. 4F). These data imply that *pha-4* activity depends on regulatory pathways operating at the *k*_*on*_ stage.

## Discussion

In this study we developed the One Click Wonder, an optimized platform for segmenting *C. elegans* embryos and nuclei. OCW provides accurate, automated segmentation across *C. elegans* developmental stages, outperforming base Cellpose models. The integration with BAAM allows robust downstream analyses: evaluation of transcriptional dynamics, dot quantification, and spatial analysis. We then applied this pipeline to examine transcriptional behavior of *pha-4*. Our data suggest that *pha-4* undergoes transcriptional bursting and revealed two distinct transcriptional populations that differ in their burst frequency rather than their burst size.

The stand-out capability of the OCW is its ability to automatically adjust model parameters for different age bins throughout embryogenesis to ensure the highest quality segmentation is always performed. As we know, embryogenesis in *C. elegans* embryos is a rapid and dynamic process. Nuclear sizes and density change drastically throughout development which generalist segmentation tools like Cellpose, Stardist, Omnipose, SAM, and Cell-ACDC are unable to cope with^1,6–9,38^. Generic models, such as those listed above, are not complete pipelines and require extensive tuning to perform tasks to the desired quality, especially across different time-stages in embryonic development. The combination of age classification with stage-specific parameters allows for automated fine-tuning of pipeline settings, ultimately reducing manual labor from days to minutes. Furthermore, the coupling of the OCW with BAAM allows for rapid post-processing and data analysis, unlocking intra-cell and intra-nuclear dynamics.

To highlight the capabilities of the OCW coupled with BAAM, we applied it to the biological case-study of *pha-4* transcription. In combination with mathematical modeling, we revealed two transcriptionally distinct *pha-4* transcription populations of cells in the embryo. These two populations are evident in the imaging to the naked eye and proves the remarkable ability of the OCW and BAAM to accurately describe transcriptional dynamics on a single-cell basis. Notably, the model that best fit the distribution of mRNA counts was one with two bursting populations of cells. As *pha-4* has been previously thought to only be expressed in pharyngeal and intestinal precursor cells, we initially thought one bursting population and one silent population of cells might best represent the data. However, occasional detections of *pha-4* transcripts in non-canonical tissues were observed, which may reflect rogue transcription events rather than tissue-specific expression patterns, and is perhaps the underlying reason why the Δ*k*_*on*_-two-population, two-state model was the best fit. Rogue transcription produces low-frequency, tissue-inappropriate transcripts which are detectable by sensitive in-situ methods but aren’t known to reflect stable, functional expression programs^39,40^. Interestingly, a recent review on transcriptional bursting by Kim et al., 2025^41^, also reported evidence for low-level, non-canonical expression events in embryonic cells using live imaging, supporting the idea that such events might be a general property of developmental transcriptional programs rather than an artifact of fixed-cell detection methods.

Additionally, we revealed that the two populations of cells, “high-expression” and “low-expression”, differed in their *k*_*on*_ values, correlating to burst frequency instead of burst size. The observed eight-fold difference aligns with documented cell-cycle-linked regulation^42^ and mirrors previous studies showing that embryonic gene regulation often modulates burst frequency^43^. This suggests that promoter activation dynamics are the primary axis of regulation, potentially coordinated with interphase timing to regulate transcription. Short cell cycles in *C. elegans* embryogenesis may favor modulation of burst frequency over size, implicating promoter-level regulation (e.g. transcription factor binding or accessibility) rather than machinery efficiency or elongation rates^44^. Live imaging studies could further test these hypotheses.

## Conclusions

The One Click Wonder provides a fast, reproducible platform for nuclear and cell segmentation, and BAAM allows for robust spatial and transcriptional quantification. Our findings highlight how combining quantitative modeling with automated image segmentation can uncover transcriptional heterogeneity in developing embryos. Together, these tools establish a modular pipeline for thorough analysis of transcriptional regulation in *C. elegans*, and can be adapted to protein or DNA analysis as well.

## Methods

### Data Acquisition

#### Nematode culture, maintenance, and growth conditions

All *C. elegans* nematode strains were maintained under standard conditions^17^. Experiments were performed with animals grown at 20°C on typical Nematode Growth Medium (NGM) agar plates that were seeded with OP50 *Escherichia coli*. All experiments were performed using N2 Bristol wild-type *C. elegans*^17^.

##### C. elegans dissection

Embryos were dissected out of adults by transferring about 50 adult *C. elegans* onto a watch glass with 1 mL MQ-H_2_O. Adults were rinsed twice with fresh MQ-H_2_O to remove residual bacteria. Using a needle, embryos were dissected under a dissecting microscope. The liquid in the watch glass containing both adult carcasses and embryos were swished to isolate embryos in the center. Dissected embryos were pipetted on to a prepared slide for smFISH.

#### smFISH experiments

smFISH experiments were performed according to Tocchini et al., 2021,^18^ with minor altercations. Probes were designed according to Tocchini et al., 2021,^18^ and ordered from IDT. Unless otherwise stated, embryos used for smFISH were dissected from hermaphrodites. The embryos were pooled in a 1.5 mL Eppindorf tube and lightly spun down using a table-top centrifuge. Embryos were then transferred to poly-L-lysine coated slides and fixative solution was applied for 5 minutes before a coverslip was placed and the slide was moved to a metal plate on dry ice. The slides were processed further according to Tocchini et al., 2021^18^. For a full list of smFISH probes, see the Supplementary table.

#### Microscopy

An Olympus SpinD CSU-W1 equipped with a Yokogawa CSU-W1 confocal scan head with a 50 µm disk, a Hamamatsu ORCA-Fusion sCMOS camera, and cellSens software was used for capturing images of smFISH-processed embryos. The UPL X APO 10x magnification was used for an overview of the slide using the DAPI channel and the individual smFISH probe channels were imaged using the UPL APO 100x magnification with a numerical aperture of 1.5. Images were processed using OMERO and assembled into figures using Adobe Illustrator software.

### One Click Wonder: A segmentation pipeline using a retrained instance of Cellpose

See supplementary information for a complete description of the Cellpose methods.

### Subsequent data analysis

#### smFISH dot counting using TrackMate

Dot detection of RNA smFISH dots was done using TrackMate (v 7.12.1), an open-source Fiji plugin designed for semi-automated detection of particles in biological images^16^. For dot detection, the Difference of Gaussians (DoG) detector was used with the dot diameter and threshold set by the user. For RNA smFISH dots, a diameter of 0.3 µm was used for all spots. The thresholds were set individually on a subset of images per dataset and then fixed for the entire dataset.

#### Spatial analysis and label application: BAAM (Biological Annotation and Association Mapper)

To determine the counts of RNA smFISH dots per embryo or per approximate cell volume, a pipeline in Python was established. The pipeline assigns each smFISH dot to the corresponding embryo or nucleus mask in which it lies. This is achieved by overlaying the dot label image onto the nuclei, dilated nuclei or embryo masks. The embryo or nucleus ID is determined by computing the mode of the pixel values at the intersection between the dot and the corresponding embryo or nucleus mask. The pipeline takes as input the original images (.vsi, .nd2, .tif(f)), the OCW embryo and nuclear label images (.tif) and the dot label images (.tif), and outputs an overview spreadsheet summarizing the number of smFISH dots per label (embryo, nuclear, dilated nuclear).

#### Statistical analysis, plot and figure generation

Plots and statistics for the One Click Wonder (OCW) pipeline were generated in Python and Jupyter Lab, using NumPy, Matplotlib, Seaborn, Pandas and SciPy^19–25^. A paired t-test was conducted on the mean values of the IOU and OCA scores from before and after retraining of the Cellpose *cyto2* model using the SciPy paired t-test function, *ttest_rel*, using the ‘less’ alternative hypothesis^25^. Statistical analyses and plots were performed in R (RStudio) using ggplot2, dplyr, tidyverse, ggtext, and ggpubr^26–32^. Log-transformations were applied where appropriate to approximate normality. Fisher’s exact test was used for unpaired categorical data with low expected frequencies; Mann-Whitney U-tests were used for non-normally distributed continuous data. Histograms and bar plots used density-scaled y-axes to account for unequal sample sizes. Schematics used public icons from bioicons.com (DBCLS and Servier). Images were processed with OMERO, and plots and figures assembled in Adobe Illustrator.

### Mathematical modelling of *pha-4* transcriptional bursting dynamics

See supplementary information for a complete description of the mathematical modelling.

## Supporting information

methodological and theoretical approaches used in the retraining of Cellpose for the One Click Wonder.

mathematical modeling

## Declarations

### Ethics approval and consent to participate

### Consent for publication

All authors have read and approved of the current version of this manuscript and agreed to its submission

### Availability of data and materials

For reviewing purposes, the images and tables are in a Zenodo dataset (this link). The codes are in a zip file in the “related files’ section, as is a README file. Or you can download from here ([this link](https://drive.google.com/drive/folders/1NomWVEPkHLp6YDm5IpSiIMWsZieS0H5p?usp=sharing)).

### Authors’ contributions

Conception and design of the One Click Wonder (T. Verheijen, A. Andriollo, S. Herbert, S.E. Mango); conception and design of BAAM and biological applications (P. Bassett, A. Angonezi, J. Chao, S.E. Mango); acquisition and analysis (P. Bassett, T. Verheijen, A. Angonezi, A. Andriollo, G. Roth); interpretation of biological data (P. Bassett, G. Roth, J. Chao, S.E. Mango); revision of the work (S.E. Mango, G. Roth, J. Chao).

## Acknowledgements

We would like to thank current and previous lab members of the Mango group and Chao group for scientific discussion. Additionally, we thank SciCORE and the IMCF for their data storage, computational resources, and internal support, WormBase for resources. Some strains were provided by the CGC, which is funded by NIH Office of Research Infrastructure Programs (P40 OD010440).

