## Supplementary material for "The One Click Wonder: a retrained automated segmentation pipeline that enables quantitative and modular analysis of *C. elegans* embryos": methodological and theoretical approaches used in the retraining of Cellpose for the One Click Wonder.

In this Supplementary Information, we describe the methodological and theoretical approaches used in the retraining of Cellpose for the One Click Wonder.

**1	Training data creation**

Label images (.tif files for which each object is given a distinct identification number) for Cellpose retraining were generated by running Cellpose on the desired image and subsequently correcting five slices in each of the three main orthogonal planes, for select embryo ages between 5 and 171 cells (Fig. 1). The labels were then manually corrected using Napari [Sofroniew et al. 2025]^1^ to best align with the raw images of embryos. This process was conducted on a set of 18 images to produce training data (Fig. 2).

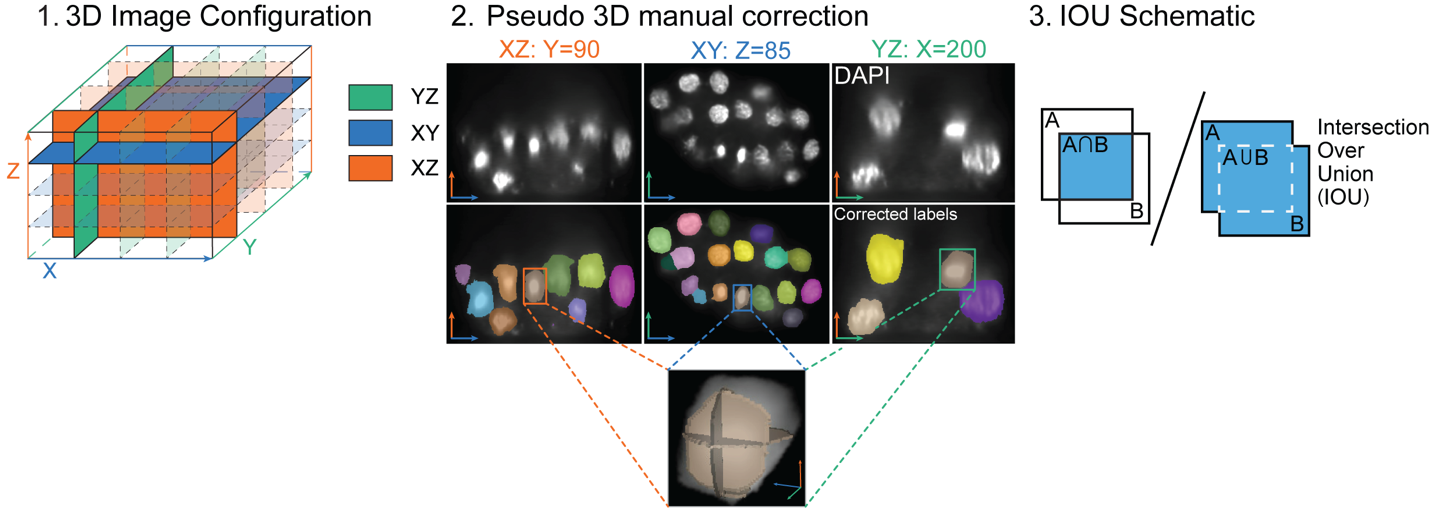

Figure 1 (1) A schematic of the representation of a 3D image where cross sections from each main axis are denoted. (2) A cross-section image from each main axis is shown with the DAPI channel and the corresponding corrected label images. For a selected nucleus, a low-resolution 3D reconstruction is provided. (3) Schematic of an intersection over union calculation between two hypothetical regions, A and B.

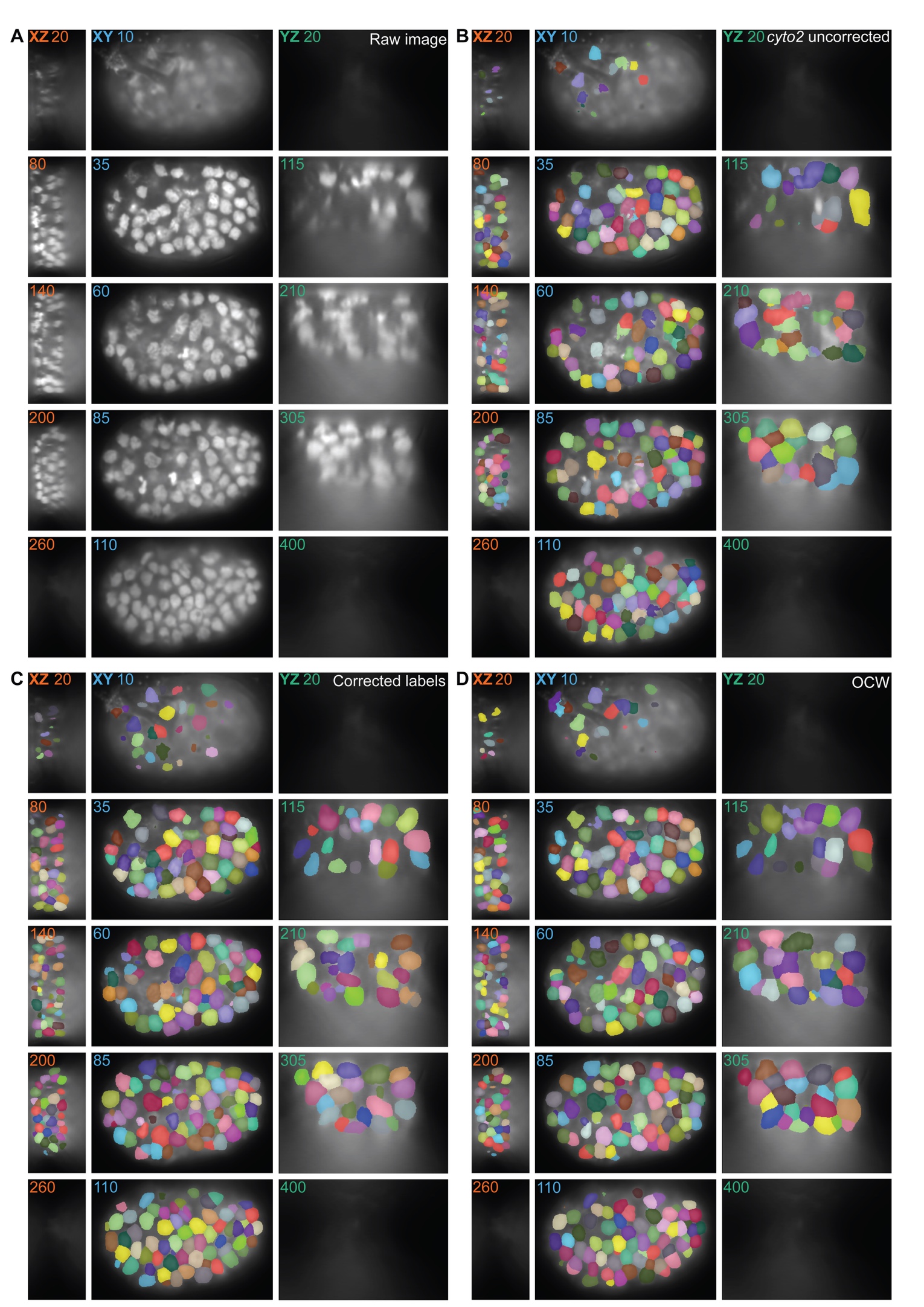

**2	Segmentation quality metrics**

Figure 2: The complete process of correcting an example embryo to include as training data. (A) Sample embryo is shown with raw DAPI channels, (B) Cellpose cyto2 uncorrected label images, (C) manual corrections of base cyto2, (D) retrained Cellpose version implemented in the OCW.

The binary intersection of the predicted and ground truth label images was computed by converting all pixels that were labeled in both images to a 1 and all pixels without a label in both images to a 0. The binary intersection was then divided by the union of both label images, which was computed by converting all pixels that had a label in at least one of the images to a 1 and all other pixels to a 0. Together, this forms intersection over union (IOU).

The Object Count Accuracy (OCA) is computed by comparing the manually counted number of nuclei to the predicted number of nuclei by the Cellpose model and rescaling the value to be between 0 and 1, where 1 represents the true number of objects. The best model is determined by the highest, equally-weighted, mean IOU and OCA score.

**3	Grid Search**

A grid search was performed before and after retraining through running Cellpose with numerous combinations of model parameters followed by computing the IOU and OCA scores on the output of the model. The best performing model is defined by the highest combined IOU and OCA score. This method was performed on different ages of *C. elegans* embryos and therefore an age-based set of optimal model parameters was selected for certain age bins (see Supplemental Table).

**4	Model Error (Loss) During Training**

The training and test error (loss) were computed and recorded during the retraining of the Cellpose model and training of the embryo age classifier at the end of each epoch. For retraining Cellpose, we used the training module of Cellpose 2.0 with a weight decay of $1\times{10}^{-4}$ and a learning rate of 0.1 with 100 epochs.^2^ For training the embryo age classifier, we used a learning rate of $1\times{10}^{-7}$ with 2500 epochs and the cross-entropy loss with an Adam optimizer [Paszke, A. et al. 2019]^3^.

**5	Cellpose**

A grid search of the cell probability threshold (“cellprob_th”), flow threshold (“flow_th”), expected object diameter (“diameter”), and minimum label size (“min_size”) parameters was performed using the IOU and OCA metrics on a retrained version of Cellpose 2.0’s *cyto2* model [Stringer et al. 2021, Pachitatiu & Stringer 2022]^5^. Common sets of parameters were found for four different age categories (cell stage, CS): 0-40 CS, 41-100 CS 101-150, and 151+ CS. The segmentation results are then saved in either a MATLAB-compatible format or as *.tif* label images depending on the user’s wishes.

Cellpose was ran on a machine with a NVIDIA RTX-A6000-12Q virtual GPU with 12 GB RAM, 48 CPU cores, and 256 GB RAM.

**6	Approximating cell volumes**

To approximate cell volumes, nuclear segmentation labels were dilated by 30-40 pixels until the labels contacted each other using the skimage.segmentation.expand_labels function.^5^ Every cell was dilated based on the nuclear shape until it met its neighboring cells or the embryo boundary as determined by the embryo segmentation labels.

**7	Imaging conditions**

The One Click Wonder is robust across a wide range of microscopes, magnifications, spatial resolutions, and embryonic developmental stages. Select parameters were optimized depending on the imaging system and dataset characteristics.

We tested the dataset using images acquired on Nikon and Olympus microscope systems:

Nikon Ti2 “seqFISH”

- NIS Elements with JOBS software
- Photometrics Prime 95B camera with 25mm field of view, F-mount, 11µm pixel size
- Lumencor SpectraX light source
- Objectives used: CFI Plan Apo Lambda 100x with numerical aperture of 1.45 with oil

Olympus SpinD and SpinSR

- Software: cellSens
- Yokogawa CSU-W1 confocal scan head (50µm disk)
- Hamamatsu ORCA-Fusion sCMOS camera
- Objectives used: UPL APO 100x with a numerical aperture of 1.5 with oil, for the SpinSR, UPL S APO 60x with a numerical aperture of 1.3 with silicone.

Table 1 Imaging conditions and optimized parameters for tested datasets. Microscope, Mode (Widefield (WF), Super Resolution (SR), Spinning Disk (SD)), Magnification (Mag.), Image format, Resolution, Percent min/max, Preprocessing, Nuclei sigma, and Oldest embryo imaged in the dataset.

| **Microscope** | **Mode** | **Mag.** | **Image format** | **Resolution (XY/Z, µm)** | **Percent min/max** | **Preprocessing** | **Nuclei sigma** | **Oldest embryo (cells)** |
| --- | --- | --- | --- | --- | --- | --- | --- | --- |
| Nikon Ti2 | WF | 100x | .nd2 | 0.11/0.20 | 60/95 | Denoising | 1.2 | ~150 |
| Olympus SpinSR | SR | 60x | .vsi | 0.034/0.13 | 60/95 | None | None | ~60 |
| Olympus SpinSR | SD | 100x | .vsi | 0.065/0.21 | 60/95 | None | None | ~100 |
| Olympus SpinD | SD | 100x | .vsi | 0.065/0.21 | 85/95 | None | 2.0 | ~500 |

Table 1 summarizes the imaging conditions and optimized parameters for the tested datasets, including the imaging mode, spatial resolution, preprocessing stems, and Gaussian blur sigma applied to images. These values represent optimized settings for the specific datasets tested and are not intended as universal defaults. For images acquired on the Nikon Ti2 widefield system, applying a denoising step followed by Gaussian blurring (sigma = 1.2) prior to running the One Click Wonder improved segmentation quality by reducing background noise and out-of-focus signal. In contrast, for images acquired on the Olympus SpinSR system, skipping the denoising and Gaussian blurring was more effective when DAPI signal appeared to be homogenously distributed across the nuclei. For images acquired on the Olympus SpinD system, DAPI signal was heterogeneously distributed across the nuclei (particularly on young embryos) and therefore, Gaussian blurring (sigma = 2.0) was effective in creating a more cohesive nuclear intensity profile, reducing the chance of over-segmentation.

**8	User guidelines**

As with all image-processing pipelines, segmentation quality depends on input data, where embryo preparation, staining quality, imaging conditions, and user-defined parameters all contribute to accurate segmentation. For optimal segmentation, we recommend high-quality nuclear staining and the optimization of imaging parameters and One Click Wonder changeable parameters. Here we present some recommendations:

Nuclear staining

- Proper washing: Embryos should be properly washed during the preparation phase to limit nonspecific binding. Specifically, when used in combination with RNA smFISH, we found that more frequent washing (instead of longer wash times) was the most effective.
- For most DAPI staining, VECTASHIELD Antifade Mounting Medium with DAPI was used and not rinsed prior to imaging.

Imaging

- Image entire embryo: For accurate age approximations using the One Click Wonder, it is important to image the entire embryo (particularly in the z-dimension) to capture the full volumes of the nuclei. Using a small z-step is also recommended.
- High quality DAPI imaging: We encourage the optimization of DAPI exposure times and laser intensity so that signal strength is strong.

Image preprocessing

- Widefield: For images captured on a Widefield system, denoising is a crucial step for removing out of focus light. For older embryos, where cells are closely packed together, this is especially important.

One Click Wonder parameters

- Gaussian blur: We recommend adding Gaussian blurring to create a more cohesive nuclear intensity profile under two circumstances:

(1) Young embryos which have relatively large nuclei and heterogenous DAPI signal strength across the nuclei
(2) DAPI signal which appears stringy and non-homogenous

- Resolution: In the pipeline, an age classifier is applied before running nuclei segmentation, where additional Cellpose parameters are inferred. The image resolution is internally used to update other size-related parameters.
