## Supplementary material for "The One Click Wonder: a retrained automated segmentation pipeline that enables quantitative and modular analysis of *C. elegans* embryos": mathematical modeling

In this Supplementary Information we describe the different mathematical models used in the study, their analysis and the fitting procedure.

#### 1 Two-state model

The two-state model of gene expression, first introduced by [Peccoud and Ycart(1995)], describes the promoter as stochastically switching between an inactive (off) and an active (on) state, with transition rates $k_{on}$ and $k_{off}$, respectively. Transcription occurs only in the on state, and RNA synthesis and degradation are modeled as Poisson processes, with rates $k_{ini}$ and $\delta$. This simplified representation treats initiation, elongation, and export as a single kinetic step.

The steady-state distribution of mRNA copy number per cell,

$$D(n) := \mathbb{P}(\text{a cell has } n \text{ mRNAs}), \tag{1}$$

has a known analytical form [Peccoud and Ycart(1995)], but evaluating it is computationally demanding due to the involvement of the confluent hypergeometric function, which lacks an efficient general-purpose numerical implementation.

To circumvent this, we compute the distribution numerically using the finite state projection (FSP) algorithm applied to the stationary chemical master equation [Gupta and Khammash(2017)]. This approach approximates the infinite state space by truncating it to a finite number of states—in our case, capping the number of mRNA molecules per cell at 150—yielding a finite system of ODEs solvable by standard methods.

Assuming the two pha-4 gene promoters act independently, the total mRNA distribution per cell is the convolution of two identical distributions:

$$D_{tot}(n) = \sum_{k \geq 0} D(k)D(n-k)$$

#### 2 Two-population, two-state model

In an embryo of 150–200 cells, approximately 25 cells are pharyngeal and intestinal precursors known to express *pha-4*. The remaining cells either do not express *pha-4*, or do so at very low levels. To capture this cellular heterogeneity, we model the embryo as a mixture of two transcriptionally distinct populations:

- The *high-pha-4-expression* population (17% of cells), corresponding to pharyngeal and intestinal precursors.
- The *low-pha-4-expression* population (83% of cells), representing the rest of the embryo.

Each population is modeled independently using the two-state model of transcription, but with distinct kinetic parameters:

- High-expression population: $k_{\text{on}}^H, k_{\text{off}}^H, k_{\text{ini}}^H, \delta$
- Low-expression population: $k_{\text{on}}^L, k_{\text{off}}^L, k_{\text{ini}}^L, \delta$

From each model, we compute the steady-state mRNA distribution, denoted $D_{\text{tot}}^H$ and $D_{\text{tot}}^L$, respectively. The total mRNA distribution observed across the embryo is then modeled as a weighted sum of the two:

$$D_{\text{embryo}}(n) = 0.17 \cdot D_{\text{tot}}^H(n) + 0.83 \cdot D_{\text{tot}}^L(n). \tag{2}$$

To identify the minimal model that best explains the observed mRNA distribution, we systematically compared a series of nested models, each varying in the number and identity of parameters allowed to differ between the two populations. The models considered, from simplest to most complex, are:

$k_{on}\Delta$**-model:** $k_{on}^H > k_{on}^L, k_{off}^H = k_{off}^L, k_{ini}^H = k_{ini}^L$

$k_{off}\Delta$**-model:** $k_{on}^H = k_{on}^L, k_{off}^H < k_{off}^L, k_{ini}^H = k_{ini}^L$

$k_{ini}\Delta$**-model:** $k_{on}^H = k_{on}^L, k_{off}^H = k_{off}^L, k_{ini}^H > k_{ini}^L$

$k_{on}k_{off}\Delta$**-model:** $k_{on}^H \neq k_{on}^L, k_{off}^H \neq k_{off}^L, k_{ini}^H = k_{ini}^L$

$k_{on}k_{ini}\Delta$**-model:** $k_{on}^H \neq k_{on}^L, k_{off}^H = k_{off}^L, k_{ini}^H \neq k_{ini}^L$

$k_{off}k_{ini}\Delta$**-model:** $k_{on}^H = k_{on}^L, k_{off}^H \neq k_{off}^L, k_{ini}^H \neq k_{ini}^L$

$k_{on}k_{off}k_{ini}\Delta$**-model:** $k_{on}^H \neq k_{on}^L, k_{off}^H \neq k_{off}^L, k_{ini}^H \neq k_{ini}^L$.

### 3 Model fitting

We compiled the mRNA counts per cell across the three replicates into a single vector $n$. Only cells from embryos containing 150–200 cells were considered, resulting in a dataset of 1775 cells.

For each version of the model, we computed the steady-state mRNA distributions of the two cell populations and combined them according to Equation (2) to obtain the total predicted mRNA distribution across the embryo, denoted $D_{\text{embryo}}$. We then calculated the log-likelihood of the observed mRNA counts:

$$LL = \sum_{i=1}^{1775} \log D_{\text{embryo}}(n_i),$$

where $n_i$ is the mRNA count in the $i$-th cell. The model parameters were estimated by maximizing this log-likelihood. For optimization, we employed a global search strategy using the `SAMIN` algorithm (Simulated Annealing) provided by the Julia package *OptimizationOptimJL*.

### 4 Model selection

We compared all versions of the model using the Bayesian Information Criterion (BIC) [Schwarz(1978)]. The BIC values for each model are listed in Table 1 and the fits are shown in Supplementary Figure XX. The model that achieved the lowest BIC, and hence the best trade-off between goodness-of-fit and model complexity, was the $k_{on}\Delta$-model in which the two population of cells only differ in their *on* rate. This result suggests that the mechanism that controls pha-4 transcription mainly controls the burst frequency of the promoter.

| model | $k^H_{on}$ | $k^H_{off}$ | $k^H_{ini}$ | $k^L_{on}$ | $k^L_{off}$ | $k^L_{ini}$ | BIC | $\Delta$BIC |
|---|---|---|---|---|---|---|---|---|
| $k_{on}\Delta$-model | 3.3 | 14206.3 | 100000 | 0.4 | 14206.3 | 100000 | 11956.3 | 0 |
| $k_{off}\Delta$-model | 0.3 | 21506.8 | 100000 | 0.3 | 4181.72 | 100000 | 12047.7 | 91.4 |
| $k_{ini}\Delta$-model | 0.3 | 100000 | 46492.5 | 0.3 | 100000 | 239117.5 | 12047.7 | 91.4 |
| $k_{on}k_{off}\Delta$-model | 1.9 | 10.5 | 145.7 | 0.4 | 22.5 | 145.7 | 11964.7 | 8.4 |
| $k_{on}k_{ini}\Delta$-model | 2.1 | 20.1 | 233.7 | 0.4 | 20.1 | 130.3 | 11965 | 8.7 |
| $k_{off}k_{ini}\Delta$-model | 0.2 | 0.03 | 2.1 | 0.2 | 5.5 | 181.3 | 12003.3 | 47 |
| $k_{on}k_{off}k_{ini}\Delta$-model | 1.6 | 6.2 | 105.2 | 0.4 | 1576.7 | 10000 | 11979.1 | 22.8 |

**Table 1:** Best fit parameter values for the different versions of the two-population, two-state model. All the rates are in $1/$mRNA lifetime unit.

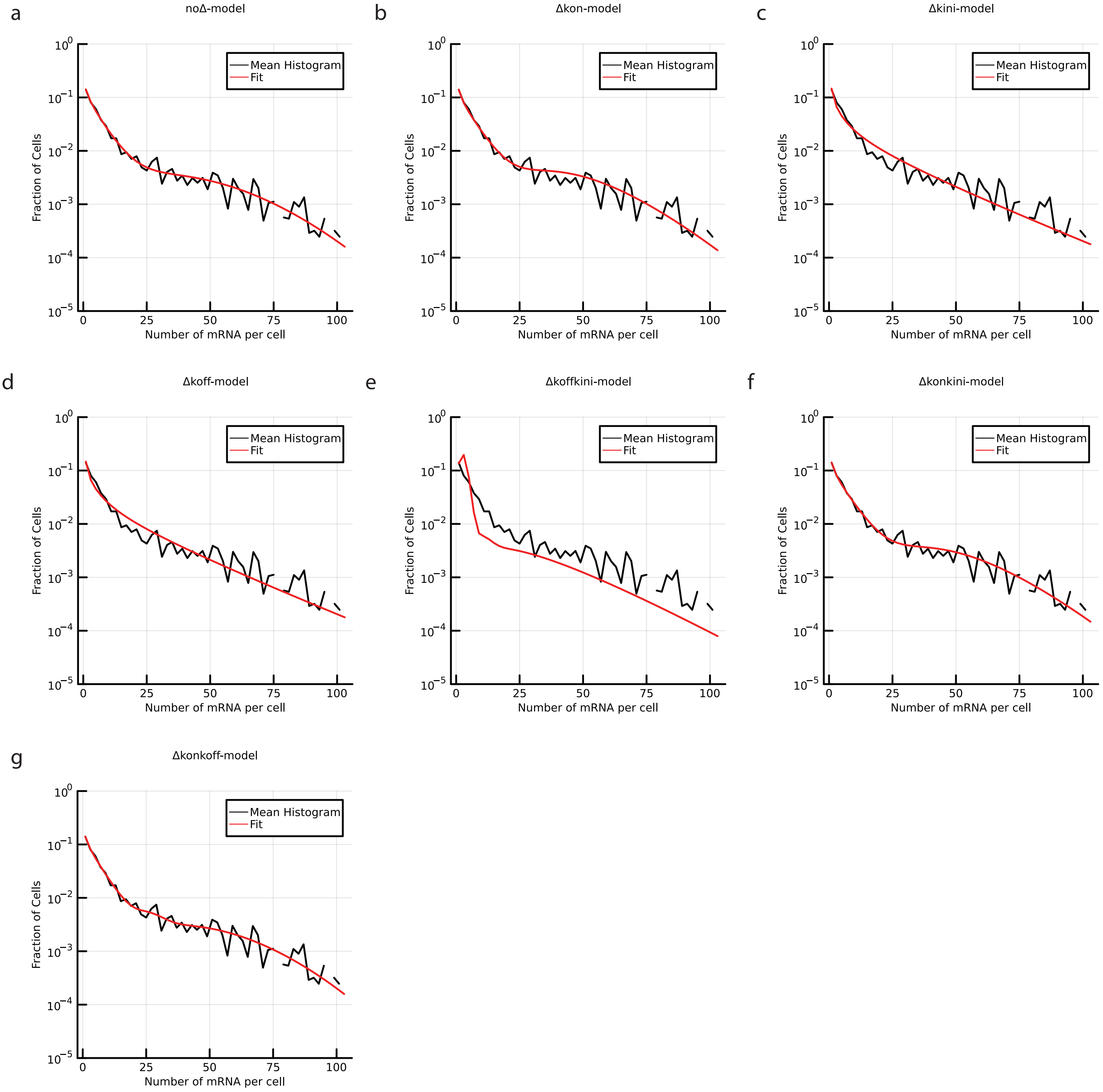
a
noΔ-model
b
Δkon-model
c
Δkini-model
d
Δkoff-model
e
Δkoffkini-model
f
Δkonkini-model
g
Δkonkoff-model
Mean Histogram
Fit
Fraction of Cells
Number of mRNA per cell
$10^0$
$10^{-1}$
$10^{-2}$
$10^{-3}$
$10^{-4}$
$10^{-5}$
0
25
50
75
100
